# Non-Canonical ERP Patterns in Second-language Sentence Processing: Evidence from Syntactic, Semantic, and Phonological Violations in Japanese Learners of English

**DOI:** 10.64898/2026.01.05.697613

**Authors:** Jananeh Shalpoush, Daniel Gallagher, Emi Yamada, Shinri Ohta

## Abstract

**Background/Objectives:** Despite notable advances in the neural mechanisms of second-language (L2) processing, few studies have systematically compared syntactic, semantic, and phonological processing of L2 within a single experimental design. We investigate the neurocognitive mechanisms underlying L2 sentence processing in native Japanese speakers with intermediate English proficiency. By integrating behavioral measures and event-related potentials (ERPs), we examined how syntactic, semantic, and phonological information influenced sentence comprehension.

**Methods:** Twenty-seven participants completed an auditory sentence judgment task involving English sentences with a syntactic, semantic, or phonological error.

**Results:** Behavioral results revealed the highest accuracy in the control and semantic conditions, while syntactic and phonological violations led to significantly lower performance, indicating greater processing difficulty in these domains. Among the three linguistic violation types, phonological violations elicited the robust ERP negativities across both time 300–500 ms and 500–800 ms time windows, while syntactic and semantic violations evoked less consistent neural responses in this L2 auditory sentence judgment task. These results suggest that mismatches in expected phonological forms hinder lexical activation, triggering a negativity that resembles an N400 but reflects different underlying processes.

**Conclusion:** We found non-canonical neural response patterns in L2 learners, characterized by sensitivity to phonological anomalies but minimal neural disruption for semantic or syntactic anomalies. The current study contributes to our understanding of L2 sentence processing in native Japanese speakers, particularly by aligning real-time neural responses with behavioral performance. This work offers implications for pedagogical practices and assessment strategies tailored to neurodiverse bilingual populations.

## 1. Introduction

The human brain is a highly dynamic and interconnected system, in which cognitive functions arise from the coordinated activity of distributed neural networks rather than isolated cortical regions [1]. Language, one of the most complex human cognitive abilities, is supported by an extensive left-lateralized fronto-temporo-parietal network that underlies both comprehension and production.

Within this network, the left inferior frontal gyrus (LIFG), which is often referred to as Broca’s area, plays a central role in morphosyntactic processing [2–13], working memory [14], and controlled aspects of language production [15]. The left superior temporal gyrus (LSTG) and adjacent temporal areas, including the superior temporal sulcus (STS), are critically involved in phonological decoding and lexical-semantic mapping [16–18]. These regions function synergistically, communicating through temporo-frontal pathways such as the arcuate fasciculus, which enables real-time integration of form, meaning, and sound. Importantly, this network interacts with the parietal and subcortical regions, forming a distributed yet functionally specialized language system capable of adapting to varying linguistic contexts and demands [19–22]. While this framework is well established for native language (L1) processing, considerably less is known about how these neural systems adapt to a second language (L2), particularly when the L1 and L2 differ strongly in linguistic typology, including morphosyntax and phonology.

Understanding how this system reorganizes during L2 learning and processing provides valuable insights into neural plasticity and the dynamic stability of linguistic representations. L2 acquisition often occurs after the period of maximal neural plasticity for language, and thus relies on partially distinct neural strategies compared to L1 processing [23–25]. While both languages engage overlapping cortical regions, L2 processing tends to recruit more extensive and bilateral networks, reflecting increased cognitive effort, working memory load, and control demands [26]. This suggests that L2 processing may not simply reuse L1 networks, but instead represents an adapted configuration shaped by proficiency, exposure, and L1-L2 structural differences. Understanding how multiple linguistic subsystems are processed in real time is therefore essential for characterizing the nature and limits of neural plasticity in adult bilinguals.

One of the most fundamental questions in L2 speech perception concerns how the brain integrates syntactic, semantic, and phonological information during the comprehension of L2 spoken sentences. These linguistic domains interact dynamically, and their interplay reflects the underlying mechanisms of linguistic representation and processing efficiency. Event-related potentials (ERPs) provide key insights into the temporal dynamics of these processes [27]. The N400, peaking around 400 ms after the presentation of a semantically anomalous word, indexes semantic expectancy and contextual integration [28]. The P600, a late positivity often emerging around 500–800 ms after the presentation of a syntactically ill-formed or ambiguous sentence, has classically been linked to syntactic reanalysis and repair [29,30], although its functional interpretation is now understood to be broader, encompassing conflict monitoring, controlled retrieval, and integration difficulty [31–33].

Multiple ERP studies on L2 learners have revealed a non-canonical response pattern where ERP components differ in amplitude, latency, or scalp distribution from those observed in native speakers [34–36]. For instance, some L2 learners may show delayed or reduced N400 effects to semantic violations, or exhibit P600-like responses to semantic rather than syntactic anomalies, a phenomenon known as the “semantic P600” [31]. Such atypical patterns are thought to reflect differences in automaticity, reliance on conscious reanalysis strategies, and influence of the L1 system on L2 processing [37,38]. These findings suggest that L2 sentence processing is characterized by reduced automaticity, increased reliance on controlled processes, and greater susceptibility to L1 transfer, especially when L1 and L2 diverge substantially.

Despite notable advances in the neural mechanisms of L2 processing, few studies have systematically compared multiple linguistic domains (e.g., syntax, semantics, and phonology) within a single experimental design.

There is a clear absence of comparable data, particularly for auditory sentence comprehension in late, unbalanced Japanese L2 learners. To date, most studies have examined these linguistic domains independently, limiting our understanding of how the brain concurrently processes and integrates multiple linguistic subsystems during real-time comprehension. Moreover, the neurocognitive mechanisms underlying how Japanese learners of English perceive and process linguistic information remain largely unexplored. Given the considerable structural and typological differences between Japanese and English, a comprehensive multimodal approach is essential.

To address these gaps, the present study investigated how late Japanese learners of English process syntactic, semantic, and phonological anomalies during auditory sentence comprehension. In particular, the study aimed to answer three central questions: (1) Do learners exhibit distinct behavioral accuracy patterns across syntactic, semantic, and phonological processing? (2) Do ERP responses to these linguistic violations elicit canonical N400 (semantic) and P600 (syntactic) effects, as typically observed in native speakers? (3) To what extent do neural responses reveal shared versus domain-specific processing mechanisms across linguistic subsystems?

Based on existing L2 ERP research, we hypothesized that semantic violations would elicit N400 effects, though potentially delayed or reduced in amplitude relative to native-like responses.

Syntactic violations were expected to evoke the P600, reflecting controlled reanalysis processes. Phonological violations were expected to elicit early-to-mid latency neural modulations, potentially cascading into semantic processing as reflected in the N400 time window. Overall, these predictions reflect canonical modulations consistent with adaptive neural reorganization in L2 processing, indicative of increased cognitive effort and reduced automaticity. However, given the typological distance between Japanese and English, we also expected ERP responses to deviate from canonical L1 patterns.

By integrating behavioral accuracy with time-resolved neural measures, this study aims to provide a multidimensional characterization of how Japanese L2 learners process multiple linguistic domains during sentence comprehension. These findings provide empirical support for the notion that L2 sentence comprehension in late Japanese bilinguals engages a qualitatively distinct neural configuration, shaped by the structural characteristics of Japanese and the processing demands of English.

## 2. Materials and Methods

### 2.1. Participants

Twenty-seven native speakers of Japanese (14 males, 13 females; age range = 18–25 years, *M* ± *SD* = 21.0 ± 2.0) participated in the study. All participants were undergraduate or graduate students at Kyushu University (Fukuoka, Japan), had normal or corrected-to-normal vision, and were self-identified as right-handed. None reported a history of neurological, speech, or hearing disorders, nor had any participant lived in an English-speaking country for more than one year.

All participants had successfully passed the mandatory, credit-bearing English language course requirement administered by Kyushu University, which typically corresponds to an intermediate level of English proficiency required for graduation. Completion of this course served as an index of participants’ formal English learning experience. Additionally, participants provided self-assessments of their English listening proficiency on a 7-point scale (M = 4.6, SD = 1.0), offering an individualized measure of their perceived auditory comprehension ability.

Informed consent was obtained from each participant after a full explanation of the experiments, in accordance with the World Medical Association’s Declaration of Helsinki. The experimental procedures were approved by the Ethics Committee of the Faculty of Humanities at Kyushu University.

### 2.2. Stimuli

The stimuli consisted of 360 English sentences, including 240 main and 120 filler sentences. The main sentences were organized into four linguistic conditions targeting common sources of difficulty for Japanese learners of English: subject-verb agreement, prepositional usage (on vs. of), countability distinctions in nouns, and definiteness and article choice (a/an/the). Within each category, sentences were distributed across four sub-conditions: syntactically correct, syntactically incorrect, phonologically incorrect, and semantically incorrect conditions (Fig. 1).

**Figure 1.**
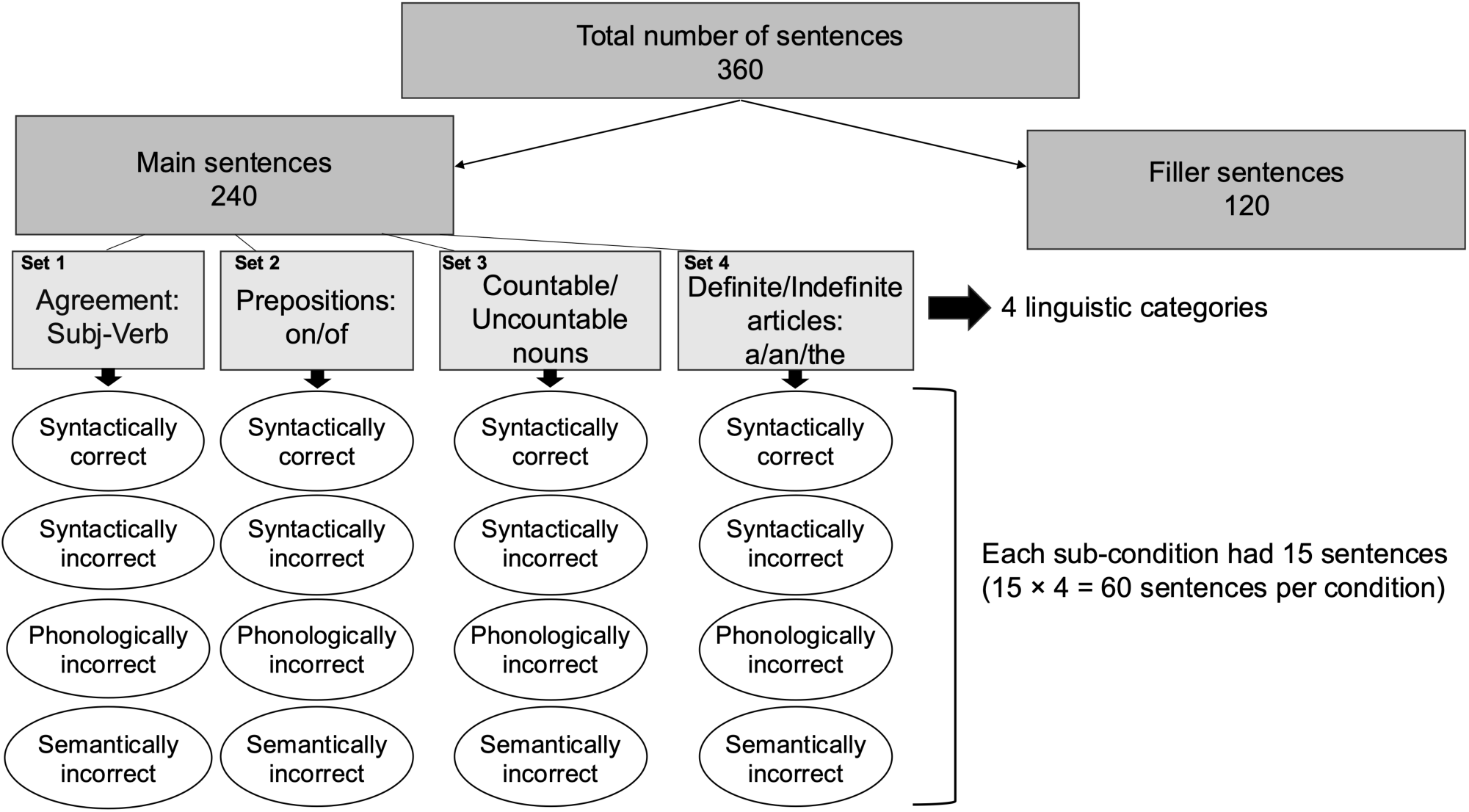
Overview of the experimental sentence design. Each of the four linguistic categories included four sub-conditions, with 15 sentences per condition (60 per category), totaling 240 main sentences and 120 filler sentences.

The syntactic violations encompassed subtypes belonging to the target domain (e.g., subject-verb agreement mismatches, omission of required prepositions, ungrammatical count/non-count usage, and omission of articles). For instance, in the subject-verb agreement set, a representative violation involved incorrect verb inflection (e.g., *The cat <u>chase</u> the mouse*), while in the prepositional set, violations omitted required prepositions (e.g., *She <u>ran out</u> time*). Countability violations involved omission of plural marking (e.g., *She saw four <u>bird</u>*), and definiteness violations omitted obligatory articles (e.g., *She visits <u>friend</u>*). This heterogeneity was intentional, as the aim was to broadly probe L2 learners’ sensitivity to different types of syntactic anomalies and to examine whether a unified ERP response pattern would emerge across structurally distinct violations. By including diverse violation types, we hoped to explore whether the neural response to syntactic anomalies in L2 learners is generalized or dissociable across subtypes. For statistical analyses, these items were collapsed into a single syntactic condition to maintain statistical power and comparability across domains. We discuss the implications of this decision in the Discussion section.

Semantic violations followed a consistent structure across all categories, replacing a critical element with an implausible yet grammatically well-formed alternative (e.g., *The cat <u>studies</u> the mouse*; *She <u>thunders</u> beautifully*; *She bought some <u>happiness</u>*; *She visits a <u>carpet</u>*). Phonological violations were also consistent across sets, created by making a controlled mispronunciation (e.g., *The cat <u>shases</u> the mouse*).

Each condition contained 15 sentences, yielding a total of 60 target sentences per category and 240 target sentences overall. The 120 filler sentences served as baseline control items and were grammatically, semantically, and phonologically well-formed and natural. All sentences were constructed by the authors to ensure precise control over lexical content, grammatical structure, and sentence length, and were designed to be easily interpretable by native Japanese speakers with intermediate English proficiency.

For auditory stimulus presentation, the sentences were synthesized using the text-to-speech software Ondoku3.com (https://ondoku3.com), ensuring a uniform speech rate, intonation, and pronunciation across all sentences, thereby minimizing acoustic variability that could confound ERP measurements. Half of the sentences (n = 180) were synthesized with a male voice, and the remaining half (n = 180) with a female voice, providing balanced gender representation across the auditory stimuli.

### 2.3. Procedure

The experiment was performed using Presentation software (Version 23.0, Neurobehavioral Systems, Inc., Berkeley, CA, www.neurobs.com), which controlled stimulus presentation, timing, and response collection.

Prior to the main experimental session, participants received task instructions in Japanese and completed a practice phase designed to familiarize them with the experimental task. Feedback was provided exclusively during this practice phase. The main experimental phase commenced after confirming, through verbal interaction, that each participant fully understood the task requirements. This approach was used to ensure that participants did not proceed to the main task until they demonstrated a solid understanding of it.

Participants then completed an auditory sentence judgment task while their brain activity was continuously recorded using electroencephalography (EEG). The task was specifically designed to assess how Japanese learners of English process distinct linguistic domains (syntactic, semantic, and phonological information) during real-time sentence comprehension. Each trial began with fixation crosses (“+++”) presented at the center of the screen for 500 ms, during which participants were instructed to maintain fixation and minimize blinking (Fig. 2). The fixation was followed by a blank screen interval of 5000 ms preceding the onset of the stimulus sequence. To indicate the type of linguistic information on which participants needed to evaluate each trial, a written cue was displayed at the beginning of every trial. The cues used in this study consisted of a single Japanese word centered on the screen: “文法” (grammar), “意味” (meaning), and “音韻” (phonology). All these cues were presented in standard-font, white-colored text without any accompanying images, shapes, or color-coding, and remained on the screen for 5000 ms before the sentence was played. Immediately following the cue, participants heard a sentence and were instructed to judge its acceptability with respect to the cued dimension by pressing one of two buttons on a response pad: the blue button to indicate “correct” and the red button to indicate “incorrect”. The auditory sentences had an average duration of approximately 3000 ms. An inter-trial interval of 200 ms was included between trials to ensure clear separation of successive events. The experiment was organized into 15 blocks of 24 trials (360 total). After each block, participants were provided with a brief rest period, during which the laboratory door was opened to reduce fatigue and maintain alertness. The entire experimental session, including task instructions, practice trials, the main task during which EEG data were recorded, and short breaks, lasted approximately 70 min.

**Figure 2.**
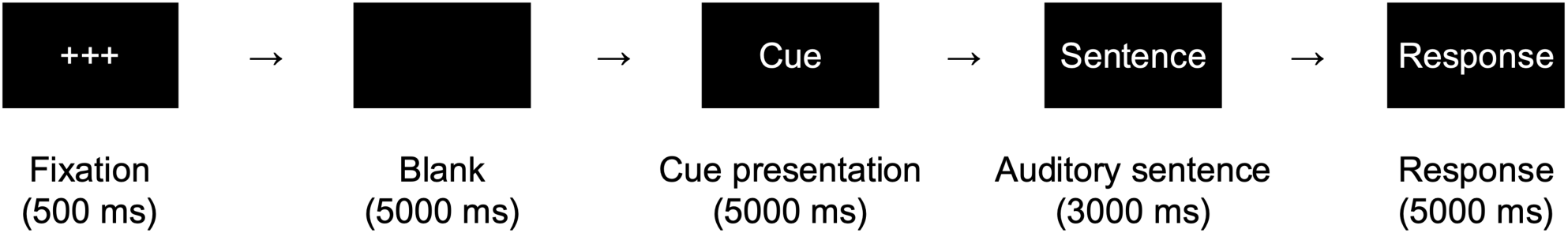
Trial sequence of a single experimental trial.

### 2.4. EEG Recording and Preprocessing

EEG data were recorded continuously using 32 Ag/AgCl electrodes positioned according to the International 10-20 system. The signal was acquired using a Neurofax EEG-1200 system (Nihon Kohden), in combination with an actiCAP electrode cap (Brain Products). Vertical and horizontal electrooculographic (EOG) activity was monitored using electrodes placed below the left eye and at the outer canthus. The electrode impedance was consistently maintained below 20 kΩ throughout the experiment. During acquisition, the EEG was sampled at 1,000 Hz. The signals were recorded relative to a specific online reference (FCz).

EEG data preprocessing and analysis were performed using MNE-Python (version 1.5.1) [39] and custom Python scripts. All procedures were performed in a Python 3.11 (64-bit) environment using the Spyder IDE (Anaconda distribution).

First, event lists were created to identify and label trigger events in the continuous EEG recordings. Following this, bad channels were visually inspected, manually rejected, and subsequently interpolated to preserve the overall channel structure. The continuous EEG data were then subjected to standard preprocessing steps, including high-pass filtering and re-referencing.

Ocular artifacts were corrected using independent component analysis (ICA), with components corresponding to blink activity identified and removed. The cleaned data were then segmented into epochs spanning from -200 ms to 2000 ms relative to the onset of the critical word. Baseline correction was applied using the pre-stimulus interval (-200 to 0 ms). Automated bad epoch rejection was performed using the autoreject algorithm to identify and exclude excessively noisy trials [40].

Grand average ERPs were computed by averaging the cleaned epochs across trials within each experimental condition for each participant. The resulting averaged data were low-pass filtered at 30.0 Hz prior to grand averaging for visualization and subsequent statistical analysis.

For ERP analyses, the N400 and P600 components were extracted. Mean amplitudes were calculated for each component within predefined regions of interest (Central, Parietal, and Centroparietal) within the established temporal windows of 300–500 ms for the N400 [27,28] and 500–800 ms for the P600 [29,30].

### 2.5. Behavioral data and ERP statistical analysis

Behavioral accuracy was calculated for each participant and condition by computing the proportion of correct responses. Mean accuracy rates were then averaged across participants and visualized using bar graphs with standard error bars. To assess whether accuracy differed statistically across conditions, a repeated-measures analysis of variance (ANOVA) was conducted with Condition (Control, Syntactic, Semantic, Phonological) as a within-subjects factor.

ERP mean amplitudes (N400 and P600) were analyzed using repeated-measures ANOVAs with factors Condition (Control, Syntactic, Semantic, Phonological) and Region (Central, Parietal, CentroParietal) for the N400 (300–500 ms) and P600 (500–800 ms) time windows. All statistical analyses were conducted in JASP (version 0.95.3, https://jasp-stats.org/). Effect sizes are reported as ω², and post-hoc comparisons were adjusted using the Holm-Bonferroni method.

## 3. Results

### 3.1. Behavioral data

Mean accuracy rates were computed for each linguistic condition to verify task engagement.

Participants exhibited the highest accuracy in the Control condition (M = 76.7 ± 2.46%), followed by the Semantic condition (M = 64.7 ± 2.39%). In contrast, performance declined substantially in the Phonological (M = 39.9 ± 3.31%) and Syntactic (M = 36.9 ± 2.52%) conditions. Values reflect mean ± standard error of the mean (SEM) (Fig. 3). For this analysis, accuracy was first computed separately for each of the four syntactic violation types and then averaged to yield a single Syntactic condition accuracy score for each participant.

**Figure 3.**
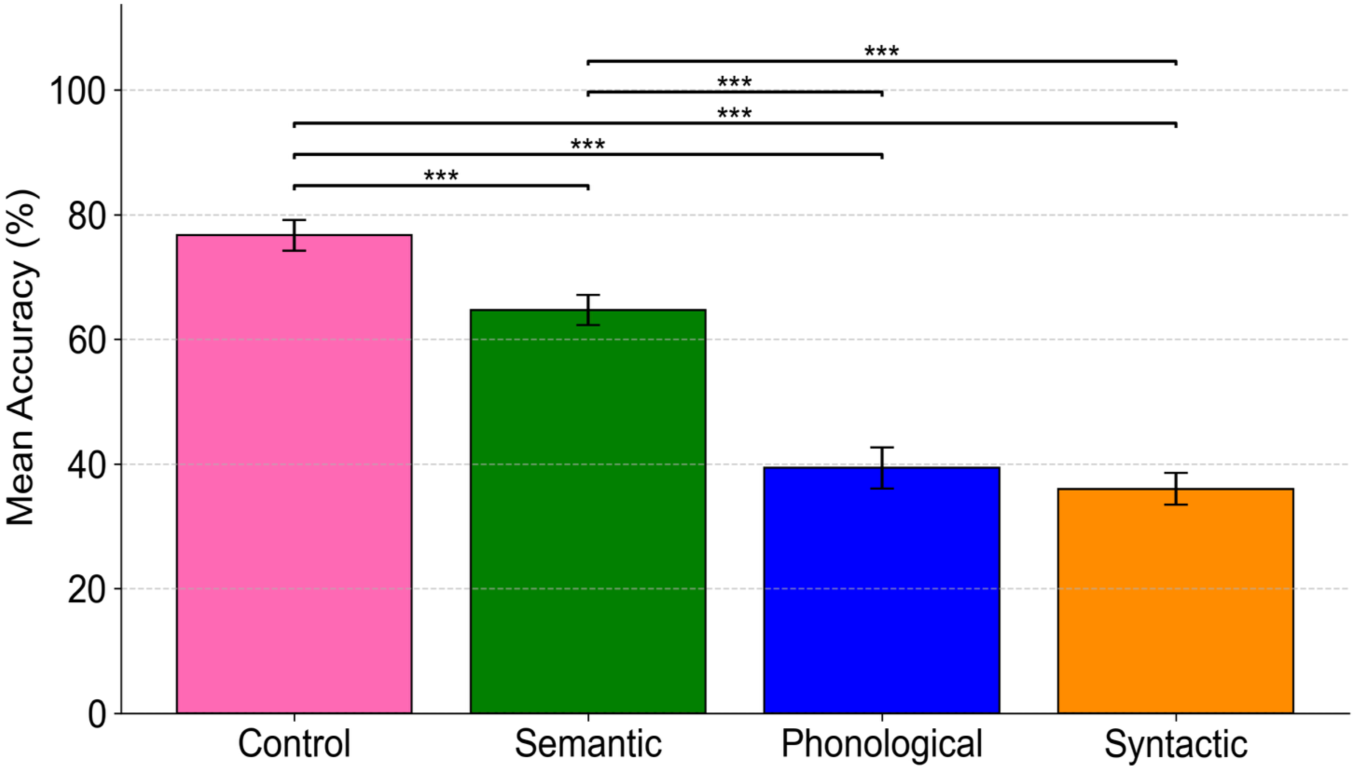
Mean accuracy rates for each linguistic condition. Error bars represent ±1 standard error of the mean (SEM). Asterisks indicate statistically significant pairwise comparisons (*p* < 0.001, Holm-Bonferroni-corrected); horizontal lines denote the tested contrasts.

To statistically evaluate the significant effect of conditions, a one-way repeated-measures ANOVA was conducted with Condition (Control, Syntactic, Semantic, and Phonological) as a within-subjects factor. The analysis revealed a significant main effect of Condition (*F*(3, 78) = 90.22, *p* < 0.001), indicating that task performance varied reliably across the linguistic conditions. Additionally, post-hoc comparisons confirmed that both Phonological and Syntactic conditions differed significantly from the Control and Semantic conditions, but not from each other (*p* > 0.10), indicating comparably low accuracy for these two violation types.

These results suggest that sentences containing syntactic violations posed the greatest difficulty, followed by those including phonological anomalies, whereas semantic violations were more readily detected. This pattern of performance aligns with previous evidence that native-language experience shapes speech perception, which can make phonological processing particularly challenging for L2 users [41–43].

### 3.2. ERP data

ERP results revealed distinct temporal and regional patterns across linguistic conditions. ERP responses were analyzed for syntactic, semantic, and phonological violations relative to control sentences in two time windows: N400 (300–500 ms) and P600 (500–800 ms) (Fig. 4).

**Figure 4.**
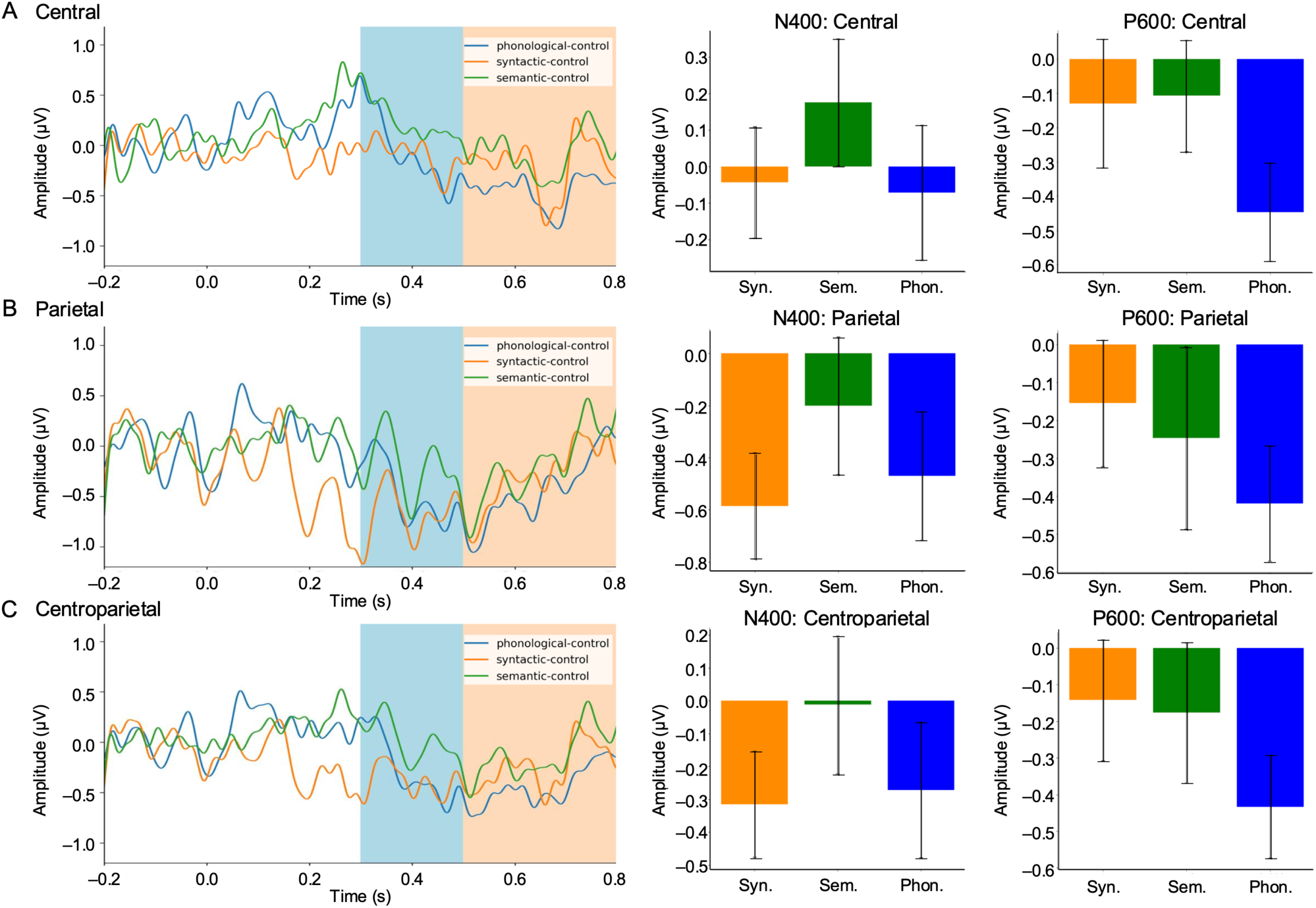
Grand-average ERP difference waves (violation minus control) for syntactic (orange), semantic (green), and phonological (blue) conditions at Central (A), Parietal (B), and Centroparietal (C) regions of interest. Shaded areas indicate the N400 (300–500 ms, blue) and P600 (500–800 ms, orange) analysis windows. Amplitude is shown in microvolts (µV), time in seconds, with negative plotted downwards. Error bars represent ±1 standard error of the mean (SEM).

As shown in Figure 4, grand-average ERP difference waves (violation minus control) are plotted for phonological, syntactic, and semantic conditions across Central, Parietal, and Centroparietal regions of interest. Since the difference wave represents the subtraction of the control condition from each violation type, deviations from zero indicate differential processing. In the 300–500 ms (N400) window, the phonological condition shows a sustained negative deflection across all regions, with the largest numerical negativity in the Parietal region (Fig. 4B). This negativity suggests increased processing demands for phonological anomalies. The syntactic condition also shows a small negative shift, but this effect is weaker and less consistent across regions (with mean amplitudes closer to zero than in the phonological condition). By contrast, the semantic condition remains near baseline across all regions, showing no clear deviation.

To statistically assess these observations, a repeated-measures ANOVA was conducted on mean N400 amplitudes during the 300–500 ms time window with factors Condition (Control, Syntax, Semantic, and Phonological) and Region (Central, Parietal, and Centroparietal) as within-subjects factors. This analysis revealed a significant main effect of Region (*F*(2,52) = 32.21, *p* < 0.001, ω² = 0.222), indicating that ERP amplitudes varied across electrode sites. No significant main effect of Condition was observed (*F*(3,78) = 1.44, *p* = 0.237), but the Condition × Region interaction was significant (*F*(6,156) = 3.10, *p* = 0.007, ω² = 0.032), suggesting that the effect of condition differed depending on scalp location.

Follow-up pairwise comparisons in the 300–500 ms time window showed a non-significant trend for the phonological condition to elicit more negative amplitudes than the control condition at parietal (*p* = 0.091) and centroparietal (*p* = 0.153) regions, with numerically larger differences posteriorly. Although the ERPs showed a broad negativity consistent with phonological sensitivity, these effects did not reach statistical significance. For the syntactic condition, a significant difference from control was observed only at the parietal region (*p* = 0.012), although the waveform lacked the typical distribution, amplitude, and shape of a canonical N400. As discussed later, this may be due to the heterogeneity of violation types and the learners’ reduced syntactic reanalysis. The semantic condition did not show significant differences from control at any region (all *ps* > 0.10), aligning with the visual pattern of minimal divergence from baseline. This lack of a canonical N400 response to semantic violations is particularly notable, given the robust N400 typically elicited by semantic anomalies in native speakers. Overall, these findings suggest that among the three linguistic violation types, phonological anomalies elicited the most robust ERP effects, particularly over posterior scalp regions. In contrast, syntactic and semantic violations showed weaker or inconsistent effects across the scalp.

In the later 500–800 ms (P600) window, there is limited evidence for a classic P600-like late positivity in response to the syntactic condition. Instead, the phonological condition continued to show negative-going activity across all regions, with the strongest effect observed over the parietal region. These patterns suggest that among the three conditions, the phonological condition elicited the most distinct ERP response relative to the control, whereas syntactic and semantic conditions elicited comparatively more subtle effects.

To statistically evaluate these observations, a repeated-measures ANOVA was conducted on mean P600 amplitudes during the 500–800 ms time window with Condition (Control, Syntactic, Semantic, and Phonological) and Region (Central, Parietal, and Centroparietal) as within-subjects factors.

We found a significant main effect of Condition (*F*(3,78) = 2.98, *p* = 0.036, ω² = .056), indicating that the ERP responses in this window differed significantly across the four conditions. However, there was no main effect of Region (*p* = 0.846), and the Condition × Region interaction was also non-significant (*p* = 0.776), suggesting that these condition effects were not strongly dependent on scalp distribution.

Follow-up pairwise comparisons indicated that the phonological condition differed significantly from the control condition at all three regions: central (*p* = 0.009), parietal (*p* = 0.030), and centroparietal (*p* = 0.016). However, these differences were driven by continued negative-going activity, not a canonical P600 activity, suggesting prolonged processing difficulty or mismatch detection rather than syntactic reanalysis.

Additionally, the syntactic condition did not differ significantly from the control condition at any region (all *ps* > 0.25), although the waveform showed a small positive shift centrally. Similarly, the semantic condition did not differ significantly from the control condition in any region (all *ps* > 0.30).

Taken together, these patterns suggest that phonological violations elicited the most robust ERP effects across both time 300–500 ms and 500–800 ms time windows, while syntactic and semantic violations evoked less consistent neural responses in this L2 auditory sentence judgment task.

Figure 5 displays the scalp topographies of the difference waves (violation minus control) for the phonological, semantic, and syntactic conditions across the Central, Parietal, and Centroparietal regions of interest during the N400 (300–500 ms) and P600 (500–800 ms) time windows. In the N400 window, the topography map shows that the phonological condition elicited a sustained posterior negativity, particularly in the parietal region, which extends into the later time window.

**Figure 5.**
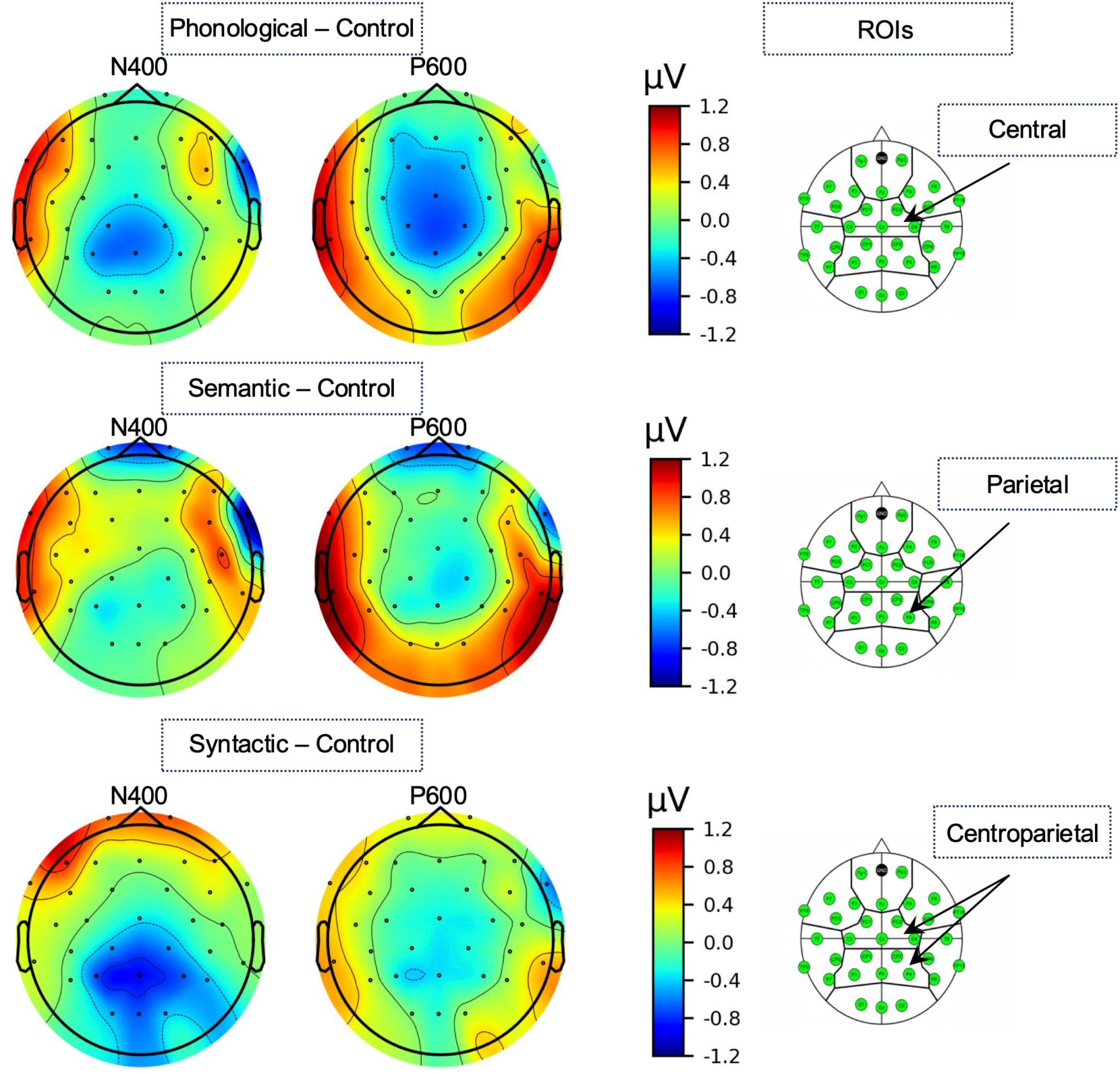
Scalp topographies of ERP difference waves (violation minus control) for the phonological, semantic, and syntactic conditions during the N400 (300–500 ms) and P600 (500–800 ms) time windows in the Central, Parietal, and Centroparietal regions of interest. These topographies support the scalp distribution described in the waveform plots, particularly highlighting the posterior negativity elicited by phonological anomalies.

This suggests increased processing effort or detection difficulty for phonological anomalies. Moreover, the syntactic condition produced potentially weaker or diffused effects across both time windows, with no clear evidence of a canonical P600. The semantic condition also showed minimal topographic modulation, with only a subtle posterior negativity in the N400 window. Overall, the topographic maps (Fig. 5) support the ERP waveforms and statistical results highlight that phonological anomalies produced the most robust ERP effects, while syntactic and semantic conditions yielded relatively inconsistent neural responses (Fig. 4).

## 4. Discussion

The present study investigated how Japanese L2 learners of English process syntactic, semantic, and phonological information in spoken L2 sentences, integrating behavioral and ERP measures to characterize both performance outcomes and underlying neural mechanisms.

Behavioral results indicated more accurate performance in the control and semantic conditions, with reduced accuracy for phonological and syntactic violations. These differences suggest that participants were less sensitive to structural and phonological anomalies, potentially reflecting reduced awareness or difficulty in detecting such violations during real-time sentence comprehension. This may indicate lower automaticity in parsing natural morphosyntax and in mapping L2 phonological forms onto lexical representations. A significant main effect of Condition confirmed these differences. We interpret this to suggest that syntactic and phonological features are less salient in L2 processing. This pattern is consistent with observations that L2 processing is characterized by persistent difficulty in automatizing grammatical features and segmenting non-native phonological input [42–44].

Unlike the behavioral data, ERP results revealed only subtle neural distinctions between linguistic conditions. In the N400 time window (300–500 ms), phonological violations elicited a small but consistent posterior negativity, particularly at the parietal region. While this negativity overlapped in timing and distribution with the canonical N400, it may not directly reflect phonological encoding. Instead, it may index downstream consequences of impaired lexical access due to earlier disruptions in phonological analysis. That is, mismatches in expected phonological forms may hinder lexical activation, triggering a negativity that resembles an N400 but reflects different underlying processes. This interpretation aligns with the notion that L2 learners, particularly at lower proficiency levels, rely more on surface-level phonological or lexical cues to process input, rather than robust form-meaning mappings.

In contrast, semantic violations did not elicit significant N400 effects relative to the control condition. The absence of a robust semantic N400, typically observed in native speakers, suggests that semantic integration processes in L2 learners may not be fully native-like. Rather than reflecting a delayed N400, this pattern may indicate qualitatively different integration mechanisms in L2 comprehension. Furthermore, task demands may have contributed to this effect. Previous work has demonstrated that the N400 amplitude is modulated by attentional allocation [45], such that reduced attention to semantic content results in attenuated or absent N400s. In the current design, the use of explicit trial-by-trial cues may have directed attention toward more perceptually salient or task-relevant features, such as phonological anomalies, limiting deeper structural or semantic reanalysis. However, since similar tasks still reliably elicit the N400 in native speakers [46,47], the present findings suggest that L2 learners may prioritize attentional resources to task-relevant features, such as anomaly detection, rather than to semantic integration. This tradeoff reveals limits in the automaticity of semantic processing and highlights the role of attentional control in L2 learners.

Although a significant parietal effect was observed for syntactic violations, the corresponding waveform lacked the characteristic shape, distribution, and amplitude of a canonical N400 or P600. This may reflect weak or distributed processing responses, possibly associated with general processing difficulty or low-level lexical disruption. One likely contributing factor is the heterogeneity of the syntactic condition. Our stimuli included subject-verb agreement errors, prepositional usage violations, countability distinctions in nouns, and definiteness and article choice errors. This diversity was a deliberate design feature intended to test whether L2 learners show converging ERP responses across different syntactic anomaly types. However, participants’ relatively low accuracy in the syntactic condition resulted in an insufficient number of correct responses per subtype. To preserve statistical power and ensure comparability with other conditions, we therefore collapsed all syntactic violations into a single condition for analysis. While this approach maintained trial counts and analytic feasibility, it may have increased within-condition variability, potentially obscuring consistent ERP effects specific to individual violation types.

Interestingly, prior research has shown that early-stage L2 learners often respond to syntactic anomalies with N400-like effects rather than P600s, suggesting reliance on lexically based detection mechanisms rather than structural reanalysis [48]. Our findings are consistent with this developmental account, where syntactic violations may be processed through disrupted lexical expectations rather than syntactic repair processes. Thus, the absence of P600 effects for syntactic violations likely reflects the limited development of syntactic reanalysis mechanisms in this learner population. At the same time, it is also possible that structural violations were not fully detected or not processed at a depth sufficient to engage repair processes. Either account offers further evidence that the neural signatures of L2 processing differ qualitatively from native-like patterns, particularly for syntactic violations.

In the later P600 window (500–800 ms), no clear P600-like positivities were observed in any condition. Phonological violations continued to elicit a sustained posterior negativity, while semantic and syntactic conditions remained close to baseline. The ANOVA revealed a significant main effect of Condition in this time window, driven primarily by the phonological condition.

Overall, the ERP results suggest that phonological violations were the most salient and consistently processed anomaly type in this study, while syntactic and semantic anomalies elicited weaker or non-canonical neural responses. These findings support the idea that L2 learners rely more heavily on phonological and lexical cues, particularly under time pressure and at lower proficiency levels. This domain-specific modulation of ERP responses is consistent with prior findings [49,50], which showed that ERP patterns in L2 learners are less native-like during early stages of acquisition, and that the nativization process varies by both linguistic domain and modality. In these ERP studies, phonological and semantic processing showed distinct developmental trajectories, suggesting that different neural mechanisms underlie the acquisition of structural versus surface-level processing skills. The current results extend this pattern by showing that phonological violations elicited clear neural effects even when syntactic and semantic anomalies did not, reinforcing the idea that multiple interacting factors, including proficiency, attention, task demands, and linguistic domain, shape L2 processing. Together, these results contribute to the growing body of evidence that L2 comprehension is not simply a delayed version of native processing, but reflects qualitatively different strategies shaped by both learner-internal and task-external constraints.

These findings underscore the importance of considering linguistic domain, task design, and learner proficiency when interpreting ERP results in L2 contexts and suggest that neural sensitivity to linguistic violations in L2 is highly selective and developmentally dynamic.

## 5. Conclusions

The findings of this study broadly support earlier works showing that L2 listeners often experience reduced ability to predict upcoming linguistic elements and weaker structural commitment during real-time comprehension (e.g., reduced N400 semantic sensitivity; attenuated P600 to syntax). These effects appear particularly for phonological anomalies, which elicited consistent neural modulation in both early and late time windows. The contrast between behavioral performance and weak neural markers suggests a dissociation between explicit decision processes and automatic neural detection in L2 comprehension [51,52]. In particular for Japanese learners, phonological mapping difficulties related to L1-L2 phonological mismatch (e.g., segmental contrasts, reduced cues to morphology) may impede form-meaning correspondence early in the processing stream, potentially decreasing the cognitive resources necessary for syntactic analysis. These findings contribute to ongoing evidence that morphosyntactic and phonological processing remain less automatized in adult-acquired languages [44,48,50], particularly in time-constrained comprehension tasks. Future work should evaluate how these domain-specific processing profiles evolve as a function of proficiency, exposure, and cognitive resources, an issue that may be especially relevant for Japanese learners given the typological characteristics of their native language [9,10,21].

## Author Contributions

Conceptualization, JS and SO; methodology, JS and DG; formal analysis, JS, DG, EY, and SO; writing—original draft preparation, JS; writing—review and editing, DG, EY, and SO; visualization, JS; funding acquisition, SO. All authors have read and agreed to the published version of the manuscript.

## Funding

This study was supported in part by JSPS KAKENHI (Grant Numbers JP24K00508, JP21K18560, JP23H05493), JST FOREST Program (Grant Number JPMJFR244J), a Research Grant from the Yoshida Foundation for the Promotion of Learning and Education, a Research Grant from the Terumo Life Science Foundation, a Research Grant from the Nakatani Foundation, a Research Grant from the Mitsubishi Foundation, and a Shimadzu Research Grant from the Shimadzu Science Foundation to Shinri Ohta.

## Institutional Review Board Statement

The study was conducted in accordance with the Declaration of Helsinki and was approved by the Ethics Committee of the Faculty of Humanities, Kyushu University (Jinbunrin-002, April 25, 2024).

## Informed Consent Statement

Written informed consent was obtained from all subjects involved in the study.

## Data Availability Statement

The data supporting the conclusions of this study are available upon request to the corresponding author. The data are not publicly available due to privacy reasons.

## Acknowledgments

The authors thank Noriko Taira for her administrative support.

## Conflicts of Interest

The authors declare no conflicts of interest.

## Notes

### Competing Interest Statement

The authors have declared no competing interest.

